# MLDSP-GUI: An alignment-free standalone tool with an interactive graphical user interface for DNA sequence comparison and analysis

**DOI:** 10.1101/745406

**Authors:** Gurjit S. Randhawa, Kathleen A. Hill, Lila Kari

## Abstract

**Summary:** MLDSP-GUI (Machine Learning with Digital Signal Processing) is an open-source, alignment-free, ultrafast, computationally lightweight, standalone software tool with an interactive Graphical User Interface (GUI) for comparison and analysis of DNA sequences. MLDSP-GUI is a general-purpose tool that can be used for a variety of applications such as taxonomic classification, disease classification, virus subtype classification, evolutionary analyses, among others.

**Availability:** MLDSP-GUI is open-source, cross-platform compatible, and is available under the terms of the Creative Commons Attribution 4.0 International license (http://creativecommons.org/licenses/by/4.0/). The executable and dataset files are available at https://sourceforge.net/projects/mldsp-gui/.

**Supplementary information:** Supplementary data are available online.

## 1 Introduction

Alignment-based methods have been successfully used for genome classification, but their use has limitations such as the need for contiguous homologous sequences, the heavy memory/time computational cost, and the dependence on *a priori* assumptions about, e.g., the gap penalty and threshold values for statistical parameters. To address these challenges, alignment-free methods have been proposed. Zielezinski *et al.*, 2017 defined two categories of alignment-free methods: those that use fixed-length word (oligomer) frequencies, and those that do not require finding fixed-length segments. MLDSP-GUI (Machine Learning with Digital Signal Processing and Graphical User Interface) combines both approaches in that it can use one-dimensional numerical representations of DNA sequences that do not require calculating *k*-mer (oligomers of length *k*) frequencies, see Randhawa *et al.*, 2019 but, in addition, it can also use *k*-mer dependent two-dimensional Chaos Game Representation (CGR) of DNA sequences, see Jeffrey, 1990; Kari *et al.*, 2015.

While alignment-free methods address some of the limitations of alignment-based methods, they still face some challenges. First, most of the existing alignment-free methods lack software implementations, which is necessary for methods to be compared on common datasets. Second, among methods that have software implementations available, the majority have been tested only on simulated sequences or on small real-world datasets. Third, the scalability issue in the form of, e.g., excessive memory overhead and execution time, still remains unsolved for large values of *k*, in the case of *k*-mer based methods.

MLDSP-GUI is a software tool that addresses all of these major challenges and introduces novel features and applications such as: An interactive graphical user interface; Output as either a 3D plot or phylogenetic tree in Newick format; Inter-cluster distance calculation; *k*-mer frequency calculation (*k* = 2, 3, 4) for analysis of under- and over-representation of oligomers; Visualisation of DNA sequences as two-dimensional CGRs; Use of Pearson Correlation Coefficient (PCC), Euclidean or Manhattan distances; Success in classifying large, real-world, datasets. The use of *k*-mer independent one-dimensional numerical representations and Discrete Fourier Transform make MLDSP-GUI ultrafast, memory-economical and scalable, while the use of supervised machine learning leads to classification accuracies over 92%. Lastly, MLDSP-GUI is user-friendly and thus ideally designed for cross-disciplinary applications.

## 2 Materials and methods

MLDSP-GUI is an interactive software tool which implements and significantly augments the ML-DSP approach proposed in Randhawa *et al.*, 2019 for the classification of genomic sequences. It is a pipeline which consists of: *(i)* Computing numerical representations of DNA sequences, *(ii)* applying Discrete Fourier Transform (DFT), *(iii)* calculating pairwise distances, and *(iv)* classifying using supervised machine learning (see Supplementary Material). More precisely, numerical representations are used to represent genomic sequences as discrete numerical sequences that can be treated as digital signals. The corresponding magnitude spectra are then obtained by applying DFT to the numerically represented sequences. A distance measure (PCC, Euclidean, or Manhattan distance) is used to calculate pairwise distances between magnitude spectra. Lastly, supervised machine learning classifiers are trained on feature vectors (consisting of the columns of the pairwise distance matrix of the training set), and then used to classify new sequences. We use 10-fold cross-validation to verify the classification accuracy. Independently, classical multidimensional scaling, see Kruskal, 1964; Karamichalis *et al.*, 2015; Solis-Reyes *et al.*, 2018, generates a visualization of the classification results in the form of a 3D Molecular Distance Map (MoDMap3D) that displays the dissimilarity-based inter-sequence relationships.

## 3 Software description

MLDSP-GUI not only gives the user the option to visualize an approximation of the inter-relationships among sequences in three-dimensional space, but also provides precise quantitative information for further analysis. The distance matrix provides the quantitative dissimilarity between any two points/sequences, while the classification accuracy scores and confusion matrix give a measure of the classification success for each individual classifier. Figure 1 shows a screenshot of MLDSP-GUI used to classify a dataset of 7,881 full mtDNA genomes of the *Flavivirus* genus. The computation of the distance matrix took 12 seconds (PCC, CGR, *k* = 6), the one-time training of the four classifiers and 10-fold cross-validation accuracy computation took 22 mins, and the classification of a new sequence 1 min.

**Fig. 1.**
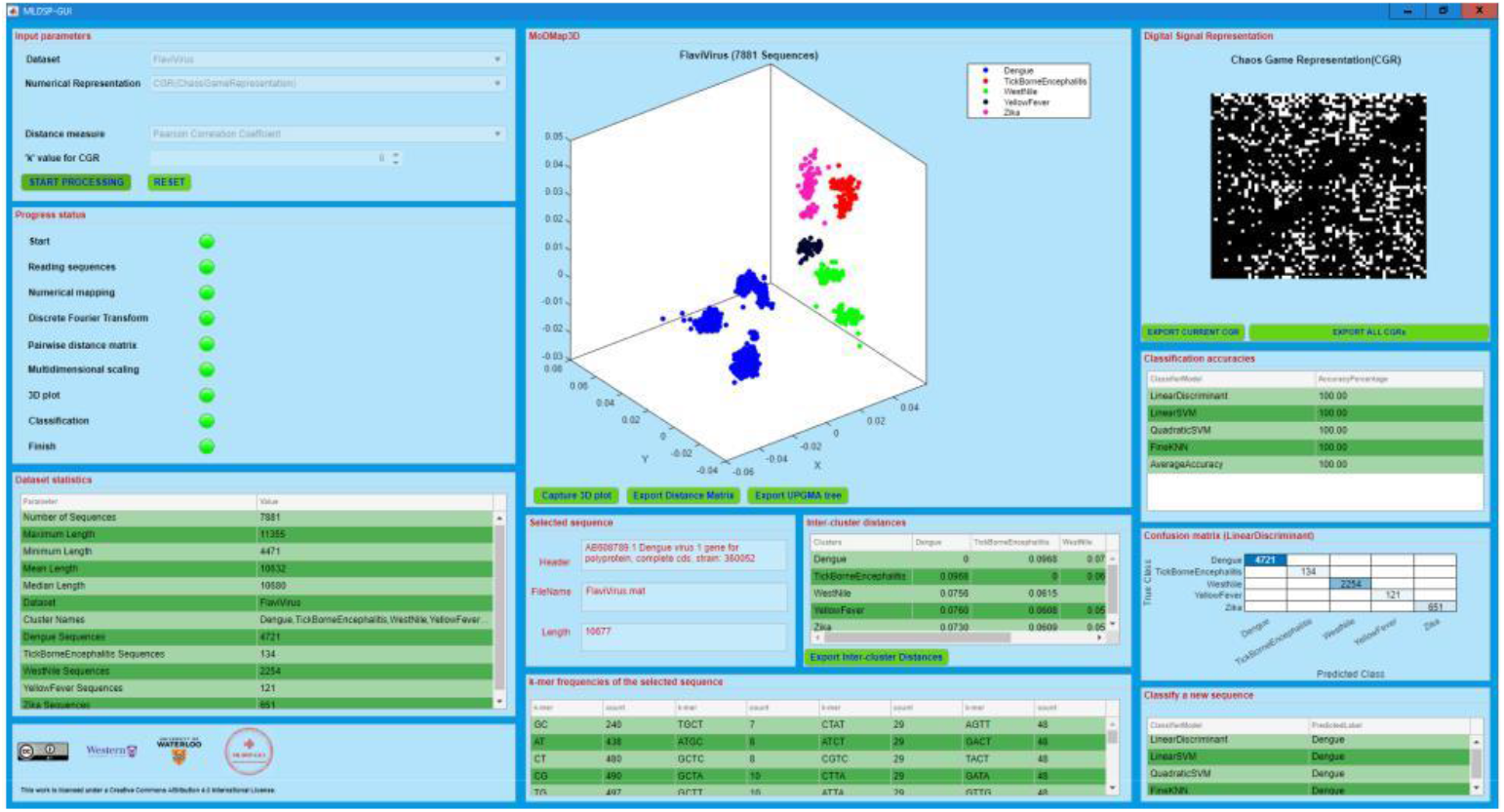
Screenshot of MLDSP-GUI showing a MoDMap3D of 7,881 full mtDNA genomes of the *Flavivirus* genus, classified into species. More details in Supplementary Material.

MLDSP-GUI takes DNA sequences in fasta file format as input. Users can select any of the provided datasets, or can input their own dataset. The tool is capable of processing a variety of DNA sequences including natural, simulated, or synthetic sequences. The 3D interactive plot can be rotated, zoomed in/out, and explored by clicking on any of the points. It auto-updates the selected point/sequence statistics such as sequence length, *k*-mer frequencies, name of parent fasta file, accession number, etc. The supervised machine learning component gives MLDSP-GUI the capability to predict the taxon of any new sequence, provided that it has been trained on a dataset containing that taxon. MLDSP-GUI is implemented using MATLAB R2019a App Designer, license no. 964054. A single executable platform-independent file is provided that can be used to install and run the software tool. The Supplementary Material file provides additional information on MLDSP-GUI features, as well as the provided datasets.

## Acknowledgements

We thank Maximillian Soltysiak for discussions, and Bianca Valente and Daniel Stueckmann for testing the software tool.

