## Supplementary material for "MLDSP-GUI: An alignment-free standalone tool with an interactive graphical user interface for DNA sequence comparison and analysis"

### MLDSP-GUI
### Supplementary Material

Gurjit S. Randhawa, Kathleen A. Hill, Lila Kari

#### A Interactive MLDSP-GUI features

MLDSP-GUI implements a four-step pipeline that takes as input a set of genomic DNA sequences and outputs their taxonomic classification. It consists of: *(i)* computing numerical representation of DNA sequences, *(ii)* applying Discrete Fourier Transform (DFT), *(iii)* calculating pairwise distances (Pearson Correlation Coefficient PCC, Euclidean, or Manhattan), and *(iv)* classifying using supervised machine learning, see Figure S1. Independently, multi-dimensional scaling uses the pairwise distance matrix to display an interactive 3D molecular distance map. The user also has an option to generate a phylogenetic tree from the pairwise distance matrix. A new sequence can be classified using the trained classifiers.

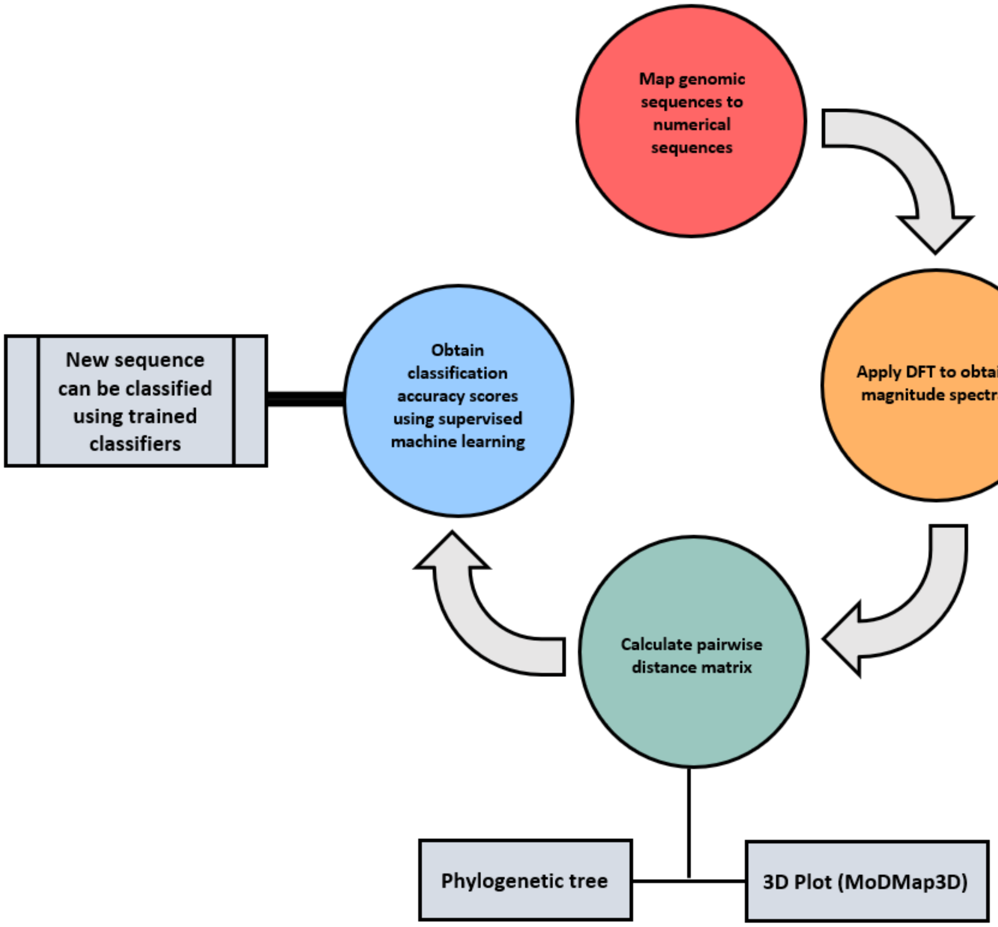


Supplementary Figure S1: MLDSP-GUI implements a four-step pipeline for data transformation from genomic sequences to taxonomic classification.

MLDSP-GUI displays results as three vertical panels, each panel subdivided into multiple sub-panel components. Figure S2 shows a test run of MLDSP-GUI on the *Flavivirus* dataset. The 7,881 complete genomes of the *Flavivirus* genus (average length 10,632 bp - the right panel shows the CGR representation of one of the *Dengue* virus genomes) are clustered into the virus species of *Dengue* (blue, 4,721 sequences), *Tick-Borne Encephalitis* (red, 134 sequences), *West Nile* (green, 2,254 sequences), *Yellow Fever* (black, 121 sequences), and *Zika* (magenta, 651 sequences). The classification accuracy using any of the four classifiers (Linear Discriminant, Linear SVM, Quadratic SVM, or Fine KNN) is 100%. MLDSP-GUI is also able to suggest classification of some virus species into subtypes, e.g., the four blue clusters correspond to the *Dengue* virus subtypes *Dengue*-1, *Dengue*-2, *Dengue*-3, and *Dengue*-4.

The next subsections of this Supplementary Material discuss the three panels (Left panel, Center panel, and Right panel) and their components in detail.

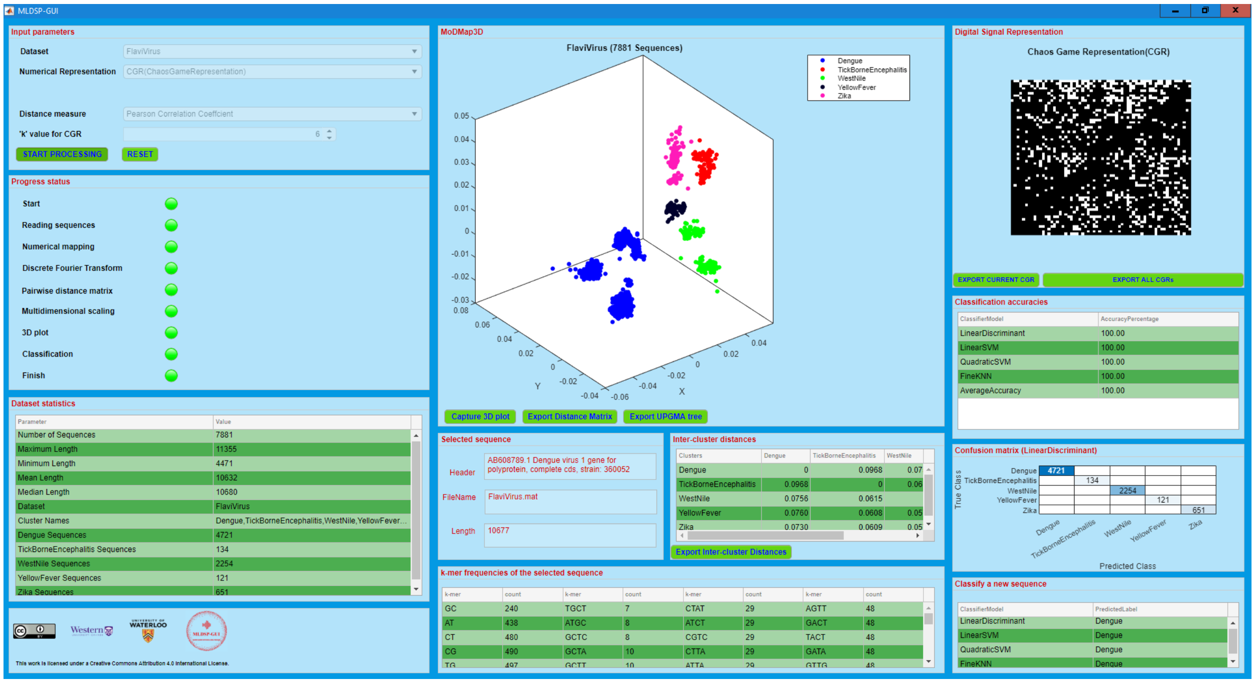


Supplementary Figure S2: MLDSP-GUI can be viewed as a combination of 3-vertical panels (Left panel, Center panel, and Right panel). Each panel has multiple sub-panel components.

All experiments were performed on an ASUS ROG G752VS computer with 4 cores (8 threads) of a 2.7GHz Intel Core i7 6820HK processor and 64GB DD4 2400MHz SDRAM.

#### A.1 Left panel

The left panel components are shown in Figure S3.

1. *Input parameters*:

   The user can select a dataset among one of the provided datasets, or "browse" to select a user-defined dataset. Some additional datasets are also provided, see Table S1.
   The user has the option to select one of the 13 one-dimensional numerical representations of DNA sequences (Integer, Integer-other variant, Real, Atomic, EIIP, purine/pyrimidine, Nearest neighbor based doublet, Codon, Just-A, Just-C, Just-G, Just-T) or the two-dimensional Chaos Game Representation (CGR).
   For example, the one-dimensional numerical representation "purine/pyrimidine" assigns A/G the value -1, and C/T the value +1, whereby the DNA sequence ACGTTAGC is represented as the numerical sequence [-1 1 -1 1 1 -1 -1 1]. If the user selects any of the one-dimensional representations, then a value for the length normalization parameter (maximum, minimum, mean or median) can be selected. The default is the length normalization using the median length.
   Alternatively, given a fixed value of the parameter $k$, the two-dimensional CGR representation of a DNA sequence simultaneously represents its $k$-mer frequencies as a two-dimensional plot (see Figure S4 for examples; for details on how to generate the CGR of a DNA sequence see Jeffrey H.J., 1990 *Nucleic Acids Res.*, *18*, 2163 – 2170). If the user selects CGR, then a $k$-value ($k$ is the length of $k$-mers to be considered when constructing the CGR) can be selected. The default value is $k = 9$ (the computations for this value could be somewhat slower), and the recommended value for a larger dataset (more than two thousand sequences) is $k = 6$.
   The user can also select a distance measure: Pearson Correlation Coefficient (PCC, the default distance), Euclidean distance, or Manhattan distance.
   After selecting the input parameters, the user can click on the Start Processing button to start the computation.
   A RESET button to reset all parameters to default is also available.

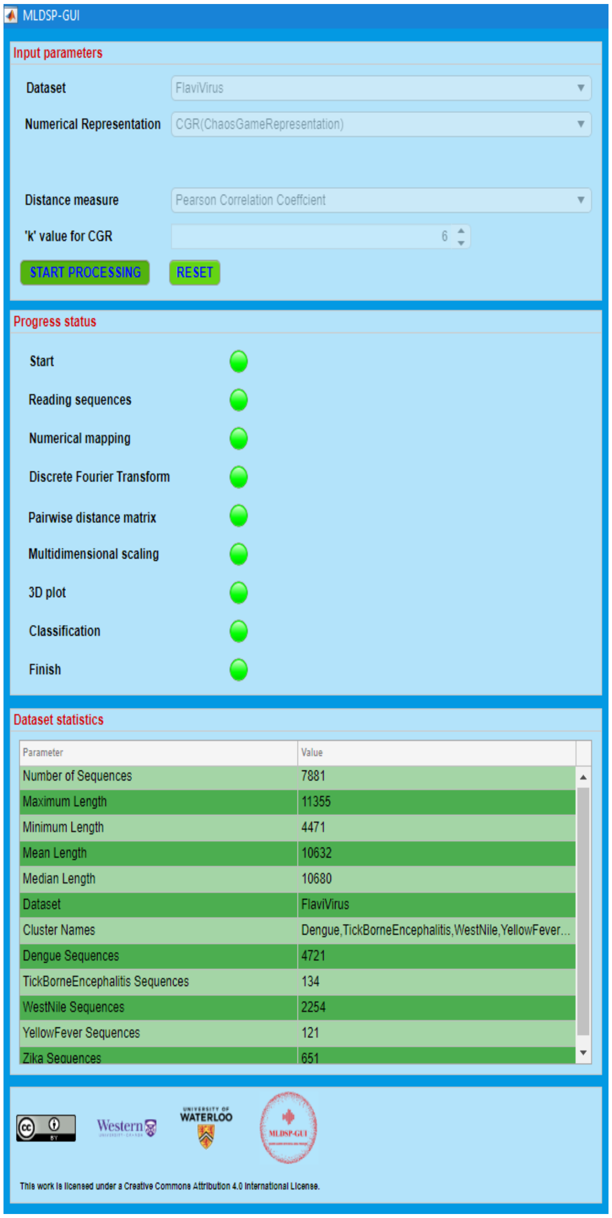


Supplementary Figure S3: Left panel components: Input parameters, progress status, dataset statistics, and logos.

2. *Progress status*:

   This sub-panel dynamically lists all the processing steps of a MLDSP-GUI computation. Each step has a colored lamp to highlight their respective status: Red means not started, yellow means in process, and green means completed.

3. *Dataset statistics*:

   This sub-panel shows some statistics of the selected dataset: number of sequences, length statistics (maximum length, minimum length, mean length, and median length), the selected dataset name, cluster names, and the size of clusters.

4. *Logos*:

   MLDSP-GUI is licensed under a Creative Commons Attribution 4.0 International License. This sub-panel contains the logos for Creative Commons, authors' affiliated institutions (The University of Western Ontario, and University of Waterloo), and MLDSP-GUI.

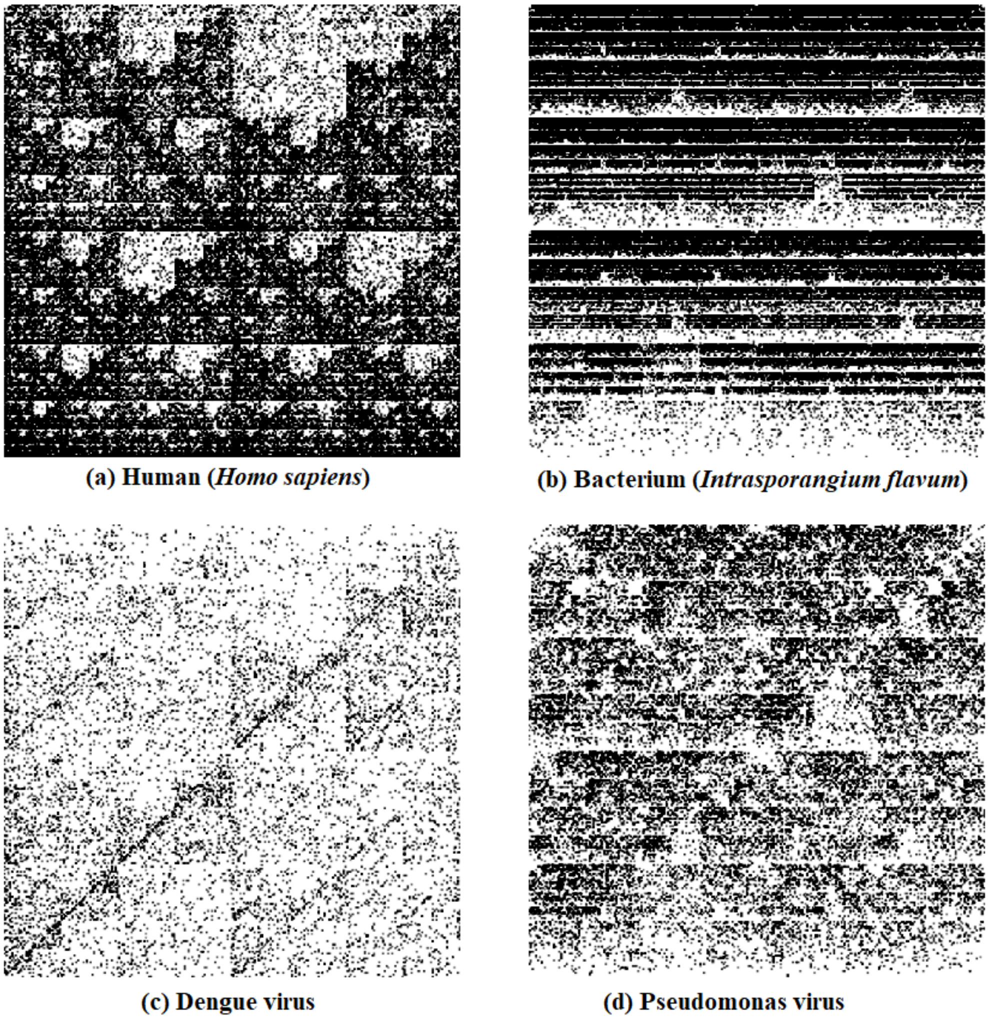


Supplementary Figure S4: Chaos Game Representation (CGR) of (a): *Homo sapiens* chromosome 1, first 100,000 bp segment, NCBI accession: $NC\_000001.11$ (b): Bacterium (*Intrasporangium flavum*) complete genome, NCBI accession: $MLJO01000003.1$ (c): *Dengue* virus 1 complete genome, NCBI accession: $AB608789.1$ (d): *Pseudomonas* phage *Andromeda* complete genome, NCBI accession: $NC\_031014.1$.

#### A.2 Center panel

The center panel components are shown in Figure S5.

1. *MoDMap3D*:

This sub-panel shows the interactive three-dimensional Molecular Distance Map (MoDMap3D) visual representation of the interrelationships among the DNA sequences in the dataset. Each point represents a DNA sequence, and the positioning of points indicates the inter-sequence relationships based on the distance used (Pearson Correlation Coefficient, Euclidean, Manhattan). Clicking on a point results in information about the selected point/sequence being displayed in the panel Selected sequence. The user also has the option to Export Distance Matrix as an excel spreadsheet, to Export UPGMA tree (UPGMA = Unweighted Pair Group Method with Arithmetic mean) in Newick phylogenetic tree format, and to Capture 3D plot of the visualized molecular distance map, as a .png file, by clicking the respective buttons.

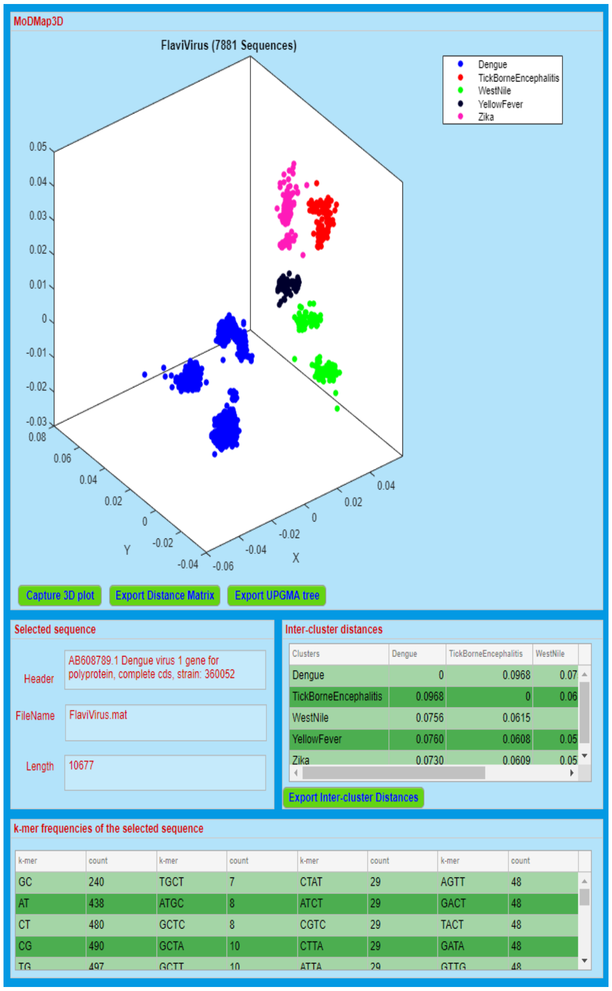


Supplementary Figure S5: Center panel components: MoDMap3D, selected sequence statistics, inter-cluster distances, and $k$-mer frequencies of the selected sequence. Export buttons for: saving 3D plot, distance matrix, UPGMA tree and inter-cluster distances.

Note that the MoDMap3D should only be viewed as a visualization tool, and is not necessarily indicative of the classification accuracy of MLDSP-GUI. This is because MoDMap3D is based on multidimensional scaling and it tries to map a multi-dimensional space onto a three-dimensional space. As such, the visual information it conveys may be imperfect (depending on the real dimensionality of the dataset that is visualized). In other words, clusters that appear to be overlapping in a MoDMap3D could in fact be perfectly separated by MLDSP-GUI, and the quantitative separability of clusters can only be accurately ascertained by looking at the accuracy scores of classifiers and at the confusion matrix.

As an example, Figure S6b shows some overlapping clusters (which indicates poor classification accuracy) in the MoDMap3D of 1,150 randomly chosen complete human mtDNA haplogroups (A, B, C, D, E, F, G, H, I, J, K, L, M, N, Q, R, T, U, V, W, X, Y, Z) sequences. However, the classification accuracy of the Linear Discriminant classifier for this dataset is reported to be 99%. The high accuracy of the quantitative classification is further confirmed by the clear visual separation obtained if we "zoom in" into the overlapping clusters of Figure S6b. Indeed Figure S6a, which displays human mtDNA haplogroups C, D, E, G, M, Q, Z, and Figure S6c which displays human mtDNA haplogroups I, K, R, W, X, both show clear separation.

As a concluding remark, when there is a discrepancy between MoDMap3D and the classification results of supervised machine learning, the latter is usually much better and also is the reliable quantitative result that should be used.

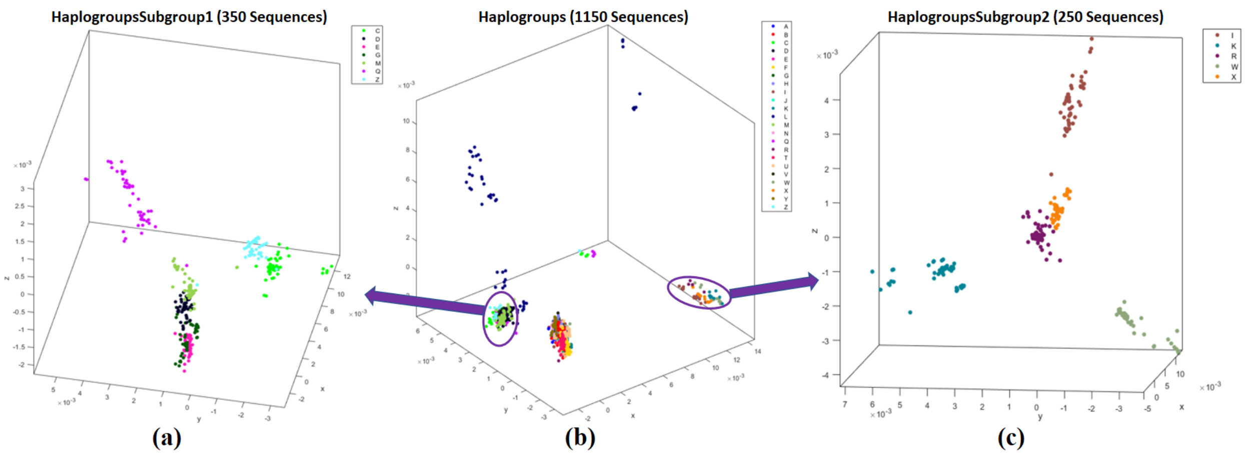


Supplementary Figure S6: "Zooming in" a ModMap3D, by re-plotting a subset of its dataset, can sometimes clarify cluster separations (separations can also be independently confirmed by the output of the supervised machine learning classifiers). Here, subfigures (a) and (c) are each obtained by re-plotting clusters which appear to be overlapping in the ModMap3D of the dataset of human mtDNA genomes from subfigure (b), as follows: **(a)** ModMap3D of 350 complete human mitochondrial genomes from the dataset in Table S1, line 13 (subset of dataset in line 12); **(b)** ModMap3D of 1,150 human mitochondrial genomes from the dataset in Table S1, line 12; **(c)** ModMap3D of 250 human mitochondrial genomes from the dataset in Table S1, line 14 (subset of dataset in line 12).

2. *Selected sequence*:

   Any point in a MoDMap3D can be selected by clicking on it. This sub-panel displays information about a selected point/sequence: Header (accession number, scientific name or other information available in the fasta file), FileName (name of its fasta file), and Length (in base pairs) of the selected sequence.

3. *Inter-cluster distances*:

   Inter-cluster distances are shown in this sub-panel. For $n$ clusters, the inter-cluster distances are shown as an $n \times n$ matrix as follows. If $M_i$ is the number of sequences in the cluster $i$, and $dist(a_s, b_t)$ gives the distance between any two sequences $a_s, b_t$, then the inter-cluster distance between any two clusters $i$ and $j$ where, $0 \leq i, j \leq n$, $1 \leq s \leq M_i$, $1 \leq t \leq M_j$, is computed as:

$$C(i,j) = \frac{\sum_{s=1}^{M_i} \sum_{t=1}^{M_j} dist(a_s, b_t)}{M_i \cdot M_j} \tag{1}$$

   The user also has the option to Export Inter-cluster Distances as an excel spreadsheet.

4. *k-mer frequencies of the selected sequence*:

   This sub-panel shows the $k$-mer frequencies (counts) for $2 \leq k \leq 4$, listed, for each $k$, in increasing order. This information can serve to analyze under-representation or over-representation of the respective oligomers.

#### A.3 Right panel

The right panel components are shown in Figure S7.

1. *Digital Signal Representation*:

   This sub-panel displays either the magnitude spectrum of the Discrete Fourier Transform applied to the numerical representation of a DNA sequence (if the one-dimensional representation was selected, Figure S8), or the CGR image of the DNA sequence (if the two-dimensional representation was selected, Figure S7).

2. *Classification accuracy*:

   The classification accuracies of six supervised machine learning classifiers (Linear Discriminant, Linear SVM, Quadratic SVM, Fine KNN, Subspace Discriminant, and Subspace KNN) using 10-fold cross validation is shown. Subspace Discriminant and Subspace KNN are omitted if the dataset has more than two thousand sequences. The average accuracy over all classifiers is also displayed.

3. *Confusion matrix*:

   A confusion matrix is displayed in this sub-panel, which changes dynamically depending on the classifier that is selected in the sub-panel above. For $m$ clusters, the $m \times m$ confusion matrix has its rows labeled by the true classes and columns labeled by the predicted classes; the cell $(i, j)$ shows the number of sequences that belong to the true class $i$, and have been predicted by the classifier to be of class $j$.

4. *Classify a new sequence*:

   MLDSP-GUI gives the option to predict the label of a new sequence, using all of the classifiers trained on a given dataset. The user can browse for a sequence (fasta file), and obtain the predicted label(s) as a result. Note that the new sequence will not be displayed in the MoDMap3D. Note also that any new sequence will be classified into one of the clusters that are displayed in the current MoDMap3D. This is an inherent limitation of supervised machine learning, in that a supervised machine learning classifier can only classify a new sequence into one of the clusters it has been trained on (it therefore classifies erroneously if the new sequence does not belong to any of the clusters that the classifier has previously "learned").

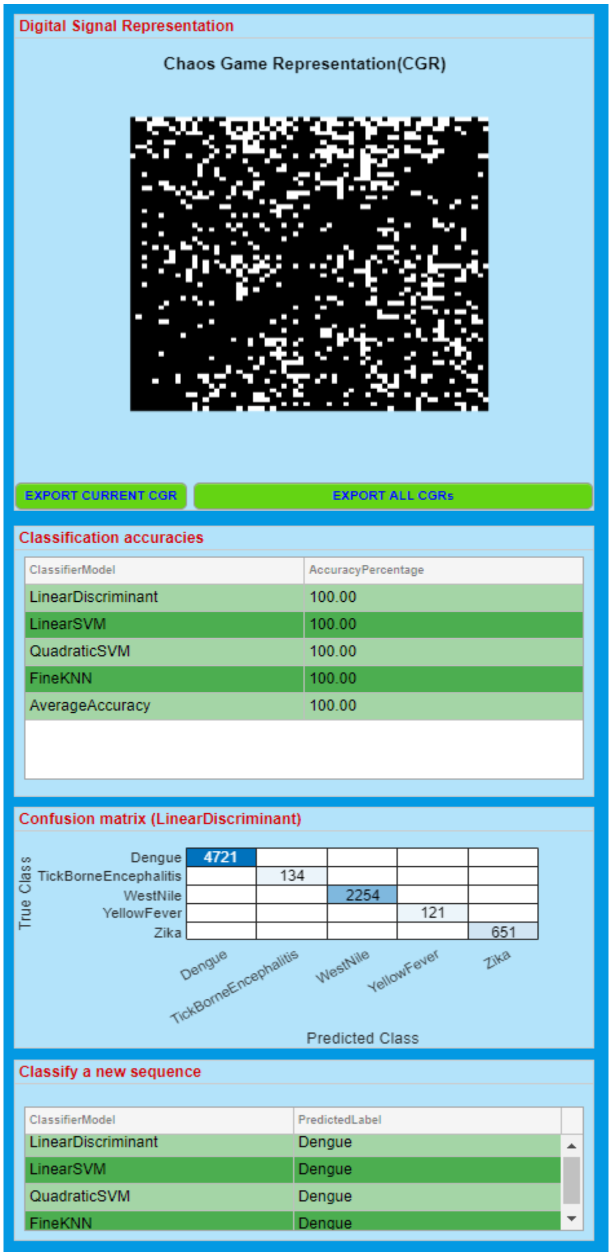


Supplementary Figure S7: Right panel components: Digital signal representation, classification accuracies, confusion matrix, and classify a new sequence.

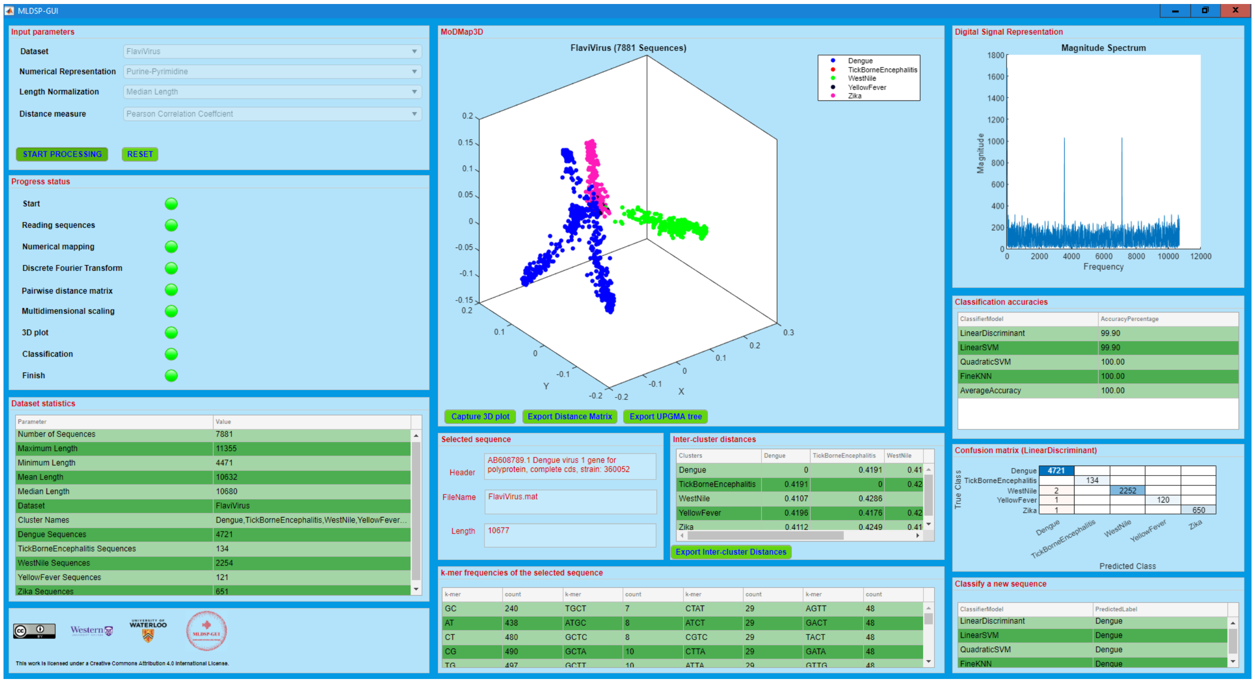


Supplementary Figure S8: MLDSP-GUI test run for the 7,881 *Flavivirus* genomes in the dataset in Table S1, line 10 using the "purine/pyrimidine" representation with length normalization to median length. The Digital Signal Representation component (top right panel) shows the magnitude spectrum of the selected point/sequence. Note that even though this is the same dataset as the one in Figure S2, the visual shape of clusters is different and the classification accuracy is lower for the Linear Discriminant classifier. The visual differences in the clusters are due to the different numerical representations used. In general, the choice of numerical representation, supervised classifier, and other parameters depend on the specific dataset, and one should choose those that achieve the best numerical classification accuracy or confusion matrix.

### B Provided datasets

Besides the datasets provided in the executable file (primates' mtDNA, influenza virus subtypes, *Flavivirus* viruses, mitochondrial disease genomes), MLDSP-GUI provides additional datasets that can be downloaded separately and imported into the already installed tool. All datasets were obtained from the NCBI Reference Sequence Database RefSeq on July 11, 2019, with the exception of the Disease-classification dataset (Table S1, line 6), which was obtained from Human Mitochondrial Database hmtDB on November 13, 2018. The additional datasets' details are given in Table S1.

Supplementary Table S1: Additional datasets provided

| S.No. | Dataset | Number of sequences | Clusters |
|---|---|---|---|
| 1 | 3classes | 3,200 | Amphibians: 264, Mammals: 1,133, Insects:1,803 |
| 2 | Amphibians | 264 | Anura: 142, Caudata: 89, Gymnophiona: 33 |
| 3 | Birds-Fish-Mammals | 4,565 | Birds (Aves): 698, Mammals (Mammalia): 2,734<br>Fish (Actinopterygii, Chondrichthyes, Coelacanthiformes, Dipnoi): 1,133 |
| 4 | ClassToSubclass (Actinopterygii) | 2,566 | Chondrostei: 28, Cladistia: 11, Neopterygii: 2,527 |
| 5 | Dengue | 4,721 | DENV-1: 2,008, DENV-2: 1,349, DENV-3: 1,010, DENV-4: 354 |
| 6 | Disease-Classification | 102 | Epilepsy: 81, Glaucoma: 21 |
| 7 | DomainToKingdom (Eukaryota) | 9,727 | Plants: 265, Animals: 8,825, Fungi: 393, Protists: 244 |
| 8 | DomainToKingdom (Eukaryota_noProtists) | 9,483 | Plants: 265, Animals: 8,825, Fungi:393 |
| 9 | FamilyToGenus (Cyprinidae) | 92 | Schizothorax: 24, Labeo: 21, Acrossocheilus: 15,<br>Acheilognathus: 11, Rhodeus: 11, Onychostoma: 10 |
| 10 | Flavivirus | 7,881 | Dengue: 4,721, TickBorneEncephalitis: 134, WestNile: 2,254, YellowFever: 121, Zika: 651 |
| 11 | Fungi | 340 | Basidiomycota: 77, Pezizomycotina: 160, Saccharomycotina: 103 |
| 12 | Human haplogroups | 1,150 | A:50, B:50, C:50, D:50, E:50, F:50, G:50, H:50, I:50, J:50, K:50, L:50,<br>M:50, N:50, Q:50, R:50, T:50, U:50, V:50, W:50, X:50, Y:50, Z:50 |
| 13 | Human haplogroups subgroup1 | 350 | C:50, D:50, E:50, G:50, M:50, Q:50, Z:50 |
| 14 | Human haplogroups subgroup2 | 250 | I:50, K:50, R:50, W:50, X:50 |
| 15 | Influenza | 38 | H1N1: 13, H2N2: 3, H5N1: 11, H7N3: 5, H7N9: 6 |
| 16 | Insects | 1636 | Coleoptera: 196, Dictyptera: 235, Diptera: 253, Hemiptera: 272,<br>Hymenoptera: 71, Lepidoptera: 442, Orthoptera: 167 |
| 17 | KingdomToPhylum (Animalia) | 8,792 | Chordata: 5,224, Cnidaria: 157, Ecdysozoa: 2,585,<br>Porifera: 64, Echinodermata: 67,<br>Lophotrochozoa: 567, Platyhelminthes: 128 |
| 18 | Mammalia | 1,075 | Xenarthrans: 36, Bats: 90, Carnivores: 145, Even-toed Ungulates: 271,<br>Insectivores: 45, Marsupials: 35, Primates: 211, Rodents and Rabbits: 242 |
| 19 | OrderToFamily (Cypriniformes) | 756 | Balitoridae: 29, Catostomidae: 14, Cobitidae: 55,<br>Cyprinidae: 597, Nemacheilidae: 61 |
| 20 | PhylumToSubphylum (Chordata) | 5,224 | Cephalochordata: 9, Craniata: 5,189, Tunicata:26 |
| 21 | Plants | 265 | Chlorophyta: 66, Streptophyta: 199 |
| 22 | Primates | 211 | Haplorrhini: 127, Strepsirrhini: 84 |
| 23 | Protists | 222 | Alveolata: 38, Rhodophyta: 80, Stramenopiles: 104 |
| 24 | SubclassToSuperorder (Neopterygii) | 1,759 | Osteoglossomorpha: 23, Elopomorpha: 63, Clupeomorpha: 92,<br>Ostariophysi: 953, Protacanthopterygii: 76, Paracanthopterygii: 48,<br>Acanthopterygii: 504 |
| 25 | SubfamilyToGenus (Acheilognathinae) | 26 | Acheilognathus: 15, Rhodeus: 11 |
| 26 | SubphylumToClass (Vertebrata) | 5,176 | Amphibians (Amphibia): 264, Birds (Aves): 698,<br>Fish (Actinopterygii, Chondrichthyes, Dipnoi, Coelacanthiformes): 2,734,<br>Mammals (Mammalia): 1,133,<br>Reptiles (Crocodylia, Sphenodontia, Squamata, Testudines): 347 |
| 27 | SuperorderToOrder (Ostariophysi) | 942 | Cypriniformes: 768, Characiformes: 40, Siluriformes: 134 |

### C Availability

MLDSP-GUI is open-source, cross-platform compatible, and is available under the terms of the Creative Commons Attribution 4.0 International license (http://creativecommons.org/licenses/by/4.0/). The executable and dataset files are available at https://sourceforge.net/projects/mldsp-gui/.
